# A human DNA methylation atlas reveals principles of cell type-specific methylation and identifies thousands of cell type-specific regulatory elements

**DOI:** 10.1101/2022.01.24.477547

**Authors:** Netanel Loyfer, Judith Magenheim, Ayelet Peretz, Gordon Cann, Joerg Bredno, Agnes Klochendler, Ilana Fox-Fisher, Sapir Shabi-Porat, Merav Hecht, Tsuria Pelet, Joshua Moss, Zeina Drawshy, Hamed Amini, Patriss Moradi, Sudharani Nagaraju, Dvora Bauman, David Shveiky, Shay Porat, Gurion Rivkin, Omer Or, Nir Hirshoren, Einat Carmon, Alon Pikarsky, Abed Khalaileh, Gideon Zamir, Ronit Grinboim, Machmud Abu Gazala, Ido Mizrahi, Noam Shussman, Amit Korach, Ori Wald, Uzi Izhar, Eldad Erez, Vladimir Yutkin, Yaacov Samet, Devorah Rotnemer Golinkin, Kirsty L. Spalding, Henrik Druid, Peter Arner, A.M. James Shapiro, Markus Grompe, Alex Aravanis, Oliver Venn, Arash Jamshidi, Ruth Shemer, Yuval Dor, Benjamin Glaser, Tommy Kaplan

**Affiliations:** School of Computer Science and Engineering, The Hebrew University of Jerusalem, Israel; Dept. of Developmental Biology and Cancer Research, Institute for Medical Research Israel-Canada, Hadassah Medical Center and Faculty of Medicine, Hebrew University of Jerusalem, Israel; GRAIL, Inc., Menlo Park, California, United States of America; Dept. of Obstetrics and Gynecology, Hadassah Medical Center and Faculty of Medicine, Hebrew University of Jerusalem, Israel; Dept. of Orthopedics, Hadassah Medical Center and Faculty of Medicine, Hebrew University of Jerusalem, Israel; Dept. of Otolaryngology, Hadassah Medical Center and Faculty of Medicine, Hebrew University of Jerusalem, Israel; Dept. of General Surgery, Hadassah Medical Center and Faculty of Medicine, Hebrew University of Jerusalem, Israel; Surgery Division, Hadassah Medical Center and Faculty of Medicine, Hebrew University of Jerusalem, Israel; Dept. of Cardiothoracic Surgery, Hadassah Medical Center and Faculty of Medicine, Hebrew University of Jerusalem, Israel; Dept. of Urology, Hadassah Medical Center and Faculty of Medicine, Hebrew University of Jerusalem, Israel; Dept. of Vascular Surgery, Shaare Zedek Medical Center, Jerusalem Israel; Dept. of Endocrinology and Metabolism, Hadassah Medical Center and Faculty of Medicine, Hebrew University of Jerusalem, Israel; Dept. of Cell and Molecular Biology, Karolinska Institutet, Stockholm, Sweden; Dept. of Oncology-Pathology, Karolinska Institutet, Stockholm, Sweden; Dept. of Forensic Medicine, The National Board of Forensic Medicine, Stockholm, Sweden; Dept. of Medicine (H7) and Karolinska University Hospital, Karolinska Institutet, Stockholm, Sweden; Dept. of Surgery and the Clinical Islet Transplant Program, University of Alberta, Edmonton, Canada; Papé Family Pediatric Research Institute, Oregon Health & Science University, Portland, OR, USA; Department of Surgery, Samson Assuta Ashdod University Hospital; Illumina, Inc., San Diego, California, United States of America

## Abstract

DNA methylation is a fundamental epigenetic mark that governs chromatin organization, cell identity, and gene expression. Here we describe a human methylome atlas, based on deep whole-genome bisulfite sequencing allowing fragment-level analysis across thousands of unique markers for 39 cell types sorted from 207 healthy tissue samples.

Replicates of the same cell-type are >99.5% identical, demonstrating robustness of cell identity programs to genetic variation and environmental perturbation. Unsupervised clustering of the atlas recapitulates key elements of tissue ontogeny, and identifies methylation patterns retained since gastrulation. Loci uniquely unmethylated in an individual cell type often reside in transcriptional enhancers and contain DNA binding sites for tissue-specific transcriptional regulators. Uniquely hyper-methylated loci are rare and are enriched for CpG islands, polycomb targets, and CTCF binding sites, suggesting a novel role in shaping cell type-specific chromatin looping. The atlas provides an essential resource for interpretation of disease-associated genetic variants, and a wealth of potential tissue-specific biomarkers for use in liquid biopsies.

**Summary paragraph:** DNA methylation, a fundamental epigenetic mark, governs chromatin organization and gene expression^1^, thus defining the molecular identity of cells and providing a window into developmental processes with wide-ranging physiologic and clinical ramifications. Current DNA methylation datasets have limitations, typically including only a fraction of methylation sites, many from cell lines that underwent massive changes in culture or from tissues containing unspecified mixtures of cells^2–6^.

We present a human methylome atlas based on deep whole-genome bisulfite sequencing of 39 sorted, primary cell types and use this dataset to address fundamental questions in developmental biology, physiology and pathology. Biological replicates are >99.5% identical, demonstrating unappreciated robustness to genetic variation and environmental perturbations. Clustering recapitulates key elements of tissue ontogeny, identifying methylation patterns retained since gastrulation. Loci uniquely unmethylated in individual cell types identify novel transcriptional enhancers and are enriched for tissue-specific transcription factors binding motifs. In contrast, loci uniquely hyper-methylated in specific cell types are rare, enriched for CpG islands and polycomb targets, and overlap CTCF binding sites, suggesting a novel role in shaping cell-type-specific chromatin looping. Finally, the atlas facilitates fragment-level deconvolution of tissue and plasma methylomes across thousands of cell-type specific regions to quantify their individual components at unprecedented resolution.

The human cell-type-specific methylation atlas provides an essential resource for studying gene regulation by defining cell-type-specific distal enhancers and regulators of 3D organization, for identifying pathological changes in DNA methylation, and for the interpretation of methylation-based liquid biopsies.

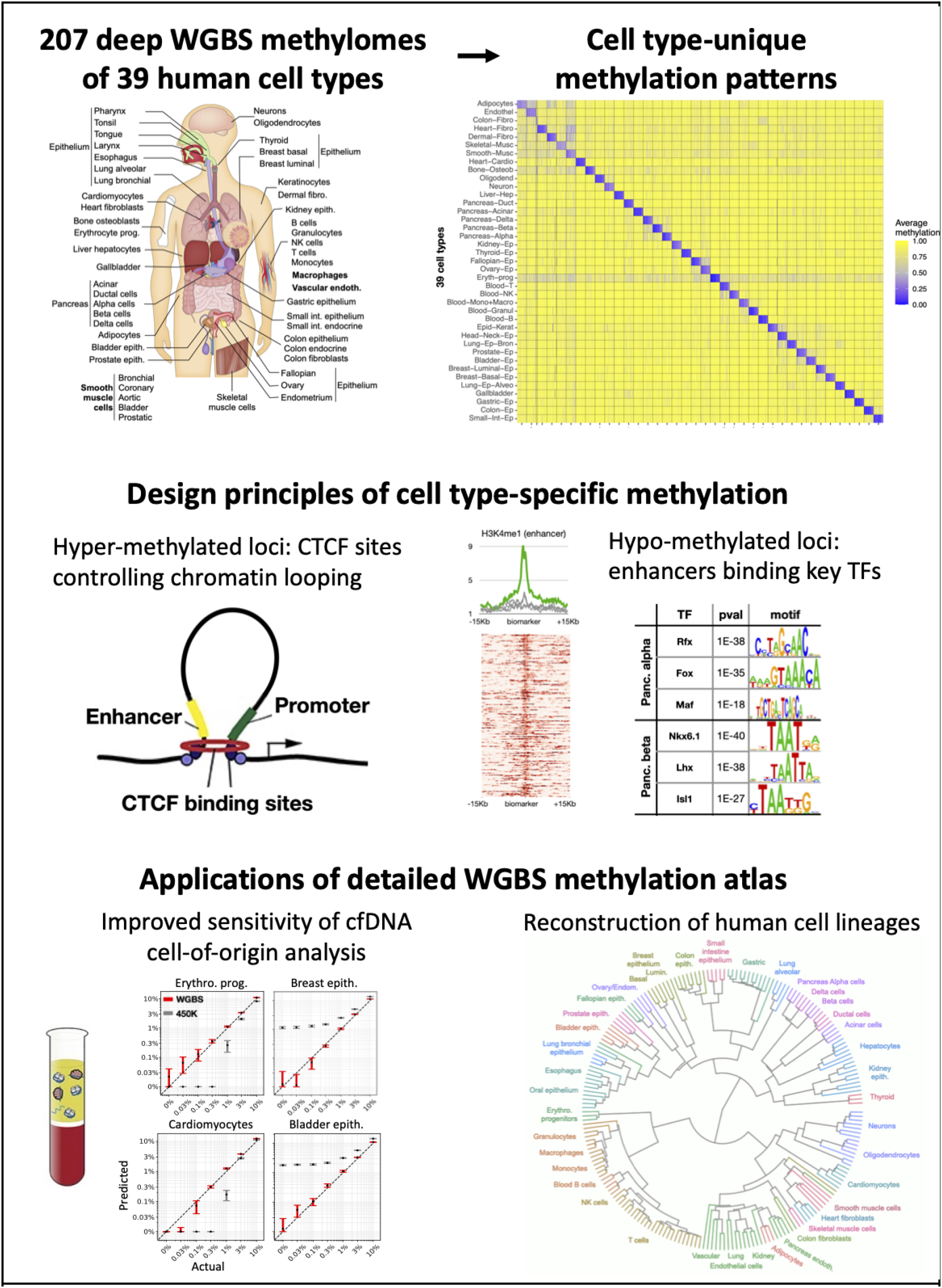

- A deep methylation atlas of 39 human cell types, sorted from healthy samples
- Methylomes record developmental history of cells
- Thousands of novel cell type-specific methylation markers
- Hypo-methylation uncovers cell type-specific regulatory map of distal enhancers
- Hyper-methylation across CTCF sites
- Cell type-specific biomarkers facilitate fragment-level deconvolution of tissues and cfDNA

## Introduction

Understanding how the same DNA sequence is interpreted differently in different cell types is a fundamental challenge of biology. Gene expression, DNA accessibility and chromatin packaging are well-established essential determinants of cellular phenotype. Underneath these lies DNA methylation, a stable epigenetic mark that underpins the lifelong maintenance of cellular identity.

Available human DNA methylation datasets suffer from major limitations. Multiple studies that characterized methylomes of embryonic development, differentiation, cancer or other settings^7–10^, have relied on the Illumina BeadChip platforms, which are limited to a predefined subset of 450K-860K CpG methylation sites, representing just 3% of the ~30 million CpG sites in the human genome^11^. Additionally, by measuring each CpG site independently, such assays overlook coordinated patterns of DNA methylation occurring in blocks, the critical functional units of DNA methylation^12,13^.

Most DNA methylation analyses interrogated primarily bulk tissue thus precluding the study of minority cell types, such as tissue resident immune cells, fibroblasts, or endothelial cells whereas others analyzed cultured cells, which may contain non-physiological methylation patterns introduced in vitro^2^. Some studies of the human methylome did analyze isolated primary cells using whole-genome bisulfite sequencing (WGBS), but their scope was limited^3,5,6^.

To overcome these limitations and to accurately characterize the human cell methylome, we performed deep genome-wide sequencing, with paired-end 150bp-long reads at an average sequencing depth of 32x (±7.2x) on FACS sorted populations of 39 human cell types obtained from freshly dissociated adult healthy tissues. We coalesced methylation patterns across the entire genome into blocks of homogeneously methylated CpG sites, and used these to study the variation of methylation patterns across cell types. Here we identify and characterize genomic regions that are differentially methylated in a tissue or cell type-specific manner, provide some vignettes of their possible biological function, and introduce a deconvolution algorithm with applications such as clinical diagnosis based on circulating cell-free DNA methylation.

## Results

### A comprehensive methylation atlas of primary human cell types

To portray genome-wide DNA methylation across a variety of cell types, we performed WGBS (150bp-long paired end reads to a mean depth of >30X) on 207 samples representing 77 primary cell types from 137 consented donors. These were mapped to the human genome (hg19) and filtered as described (Methods, Extended Table 1 and Fig. S1).

Cell types analyzed (Figure 1) represent most major human cell types, allowing a composite view of physiological systems (e.g. GI tract, hematopoietic cells, pancreas), as well as a comparison of similar cell types in different environments (e.g. tissue-resident macrophages).

**Figure 1.**
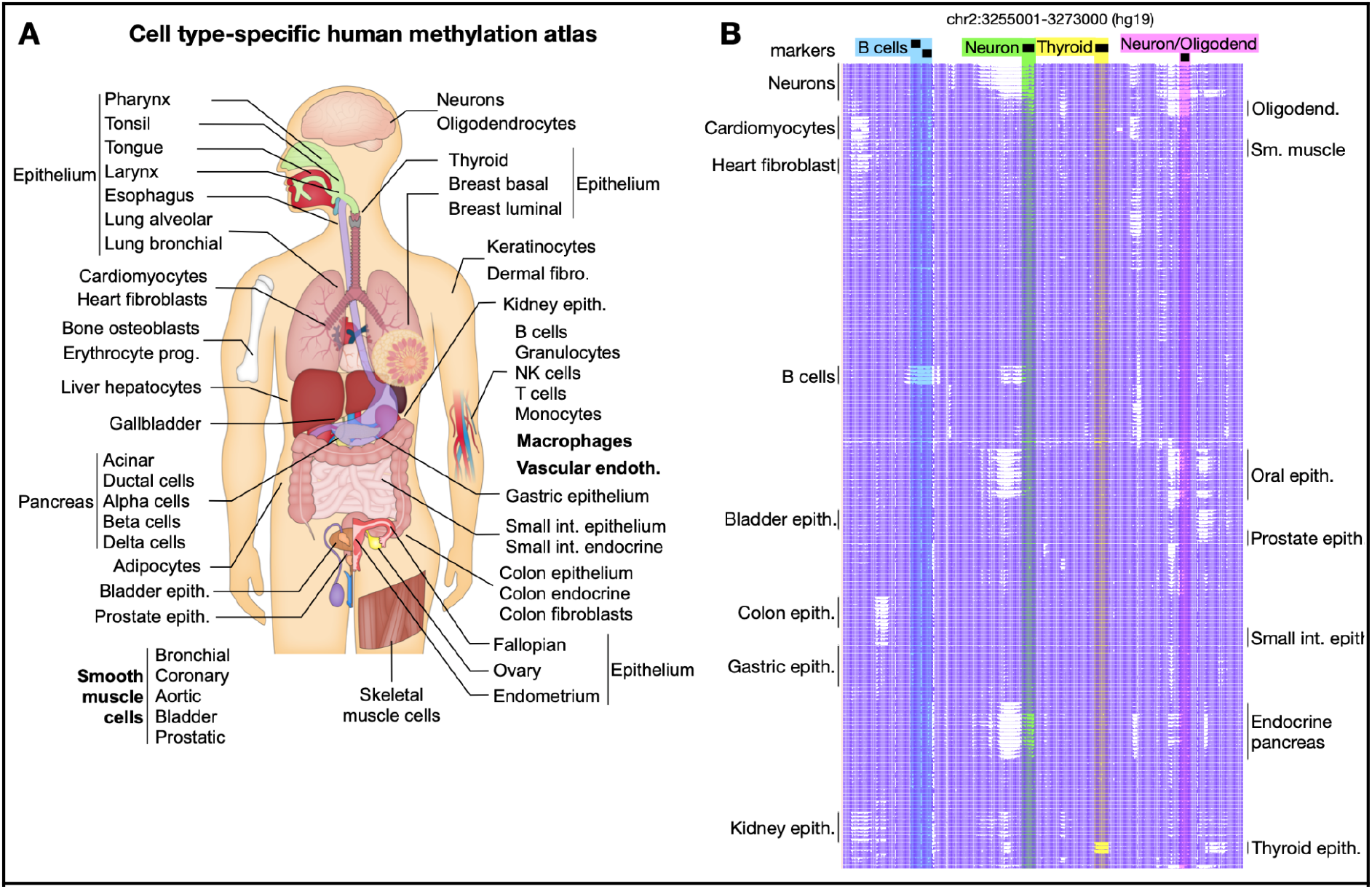
Methylation atlas of the adult human body. **(A)** 207 healthy samples were obtained from adult humans, isolated and deeply sequenced (WGBS, mean depth >30x), to form a comprehensive human cell type-specific methylation atlas. **(B)** DNA methylation patterns of 207 methylomes (rows) across 344 CpG sites (columns) within a 18Kb region. Highlighted are regions differentially unmethylated in B cells (blue), neurons (green), thyroid epithelium (yellow) and neurons/oligodendrocytes (pink).

### Identification of human methylation blocks

The 207 methylomes show great similarities between replicates, with distinctive changes between cell types in a block-like manner as shown in Fig. 1. We sought to identify genomic regions that are differentially methylated in specific cell types, to shed light on cell type-specific biological processes, define cell identity, and facilitate development of methylation biomarkers to identify the cellular origin of circulating cfDNA fragments^1,12–19^.

We developed *wgbstools*, a computational machine learning suite to represent, compress, visualize and analyze WGBS data (https://github.com/nloyfer/wgbs_tools). We segmented the genome into 7,264,350 non-overlapping continuous blocks, by identifying changepoints in DNA methylation patterns across multiple conditions. Each block spans highly correlated CpG sites that are similarly methylated in each sample, but may co-vary across cell types. We retained 2,807,024 methylation blocks of ≥3 CpGs, with an average length of 532bp (IQR=551bp) and 8 CpGs (IQR=5 CpGs, Extended Figure S2). These compact genomic units are more straightforward to robustly analyze than individual CpG sites, and due to the regional nature of methylation can be viewed as the biological “atoms” of human DNA methylation13.

### The extent of inter-individual variation in cell type methylation

Methylation patterns were extremely robust across different individuals. For most cell types, ≤0.5% of blocks show a difference of 50% or more across different donors, compared to 4.9% among samples of different cell types (Extended Figure S3). This high similarity in DNA methylation across donors is on par with the estimated inter-individual variability of genomic sequence^20^. While the definition of 50% is somewhat arbitrary, other thresholds (35%-50%) show a similar trend, with ≤0.5% variable blocks. Similar inter-individual variation was observed in replicates obtained from different laboratories (Table S1). Strikingly, for cell types with n≥3 biological replicates, 200/201 samples (99.5%) showed the highest similarity to another replicate (rather than another cell type from the same donor). These results demonstrate the reproducibility of preparations, but also highlight a fundamental biological phenomenon: determination of the epigenome primarily by cell type-specific programs, rather than genetic or environmental factors.

### Methylation patterns record human developmental history

Whereas DNA methylation patterns reflect the functional identity of a cell, they could also be used to track its developmental history. To identify patterns shared by the progeny of early progenitors, we calculated the average methylation within blocks of ≥4 CpGs, and selected those showing the highest variability across all samples (21K blocks, top 1%, Extended Table S2 and Figures S3,S4). We then clustered all 207 methylomes using an unsupervised agglomerative algorithm (UPGMA) that iteratively identifies and connects the two closest samples, regardless of their labeling^21^. This analysis systematically grouped together biological replicates of the same cell type (Figure 2), supporting reproducibility of cell isolation and suggesting that 3-4 replicates of each normal cell type are sufficient to infer its methylation patterns for practical applications such as biomarker identification.

**Figure 2:**
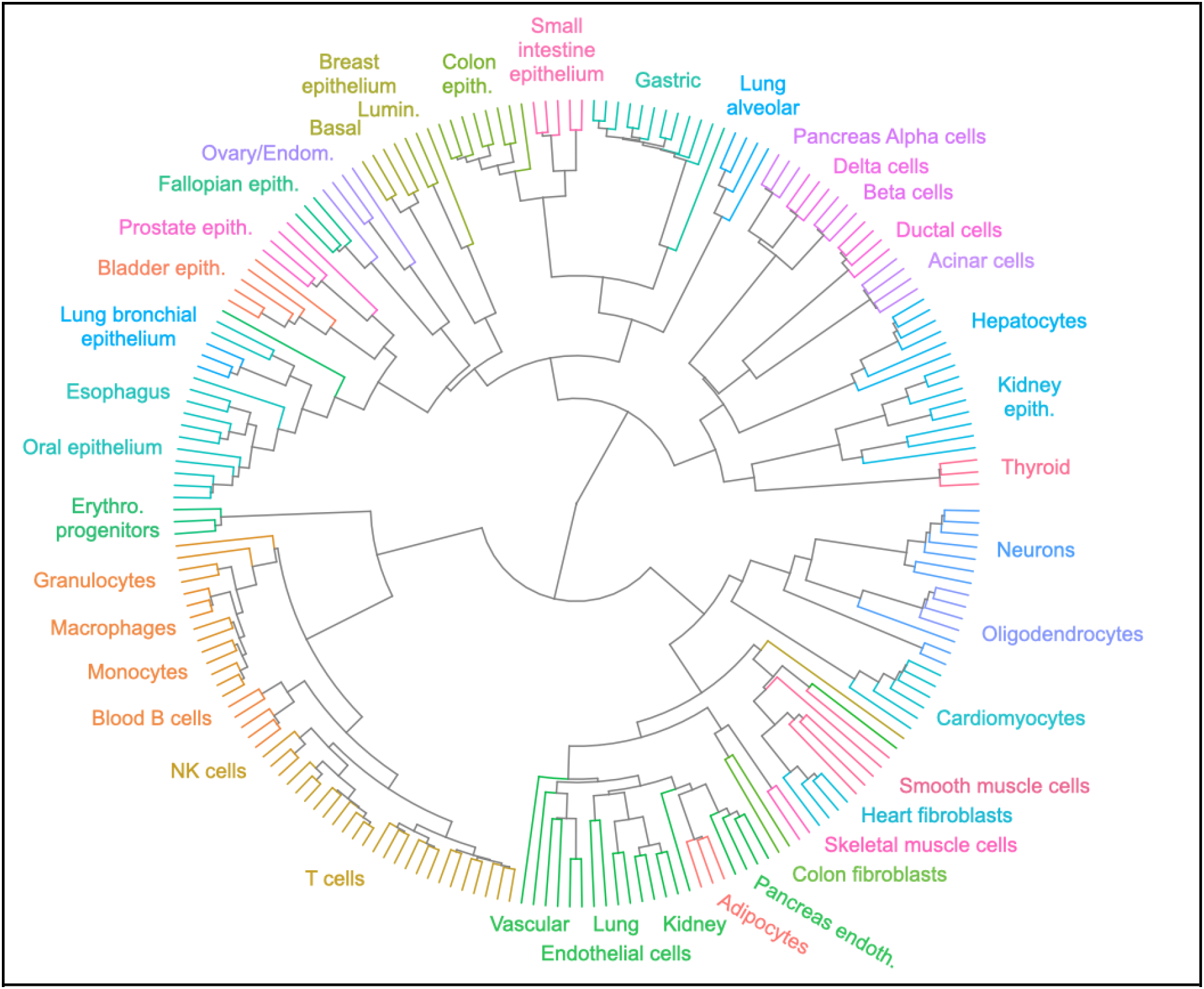
Unsupervised agglomerative clustering reflects human developmental lineage of healthy cell types. Cell types are indicated by edge colors.

Strikingly, the resulting fanning diagram recapitulates key elements of lineage relationships among human tissues. For example, pancreatic islet cell types (alpha, beta, delta), which originate from the same embryonic endocrine progenitor^22^, densely cluster together. Consistently with methylomes reflecting lineage rather than function, islet cells further cluster with pancreatic duct and acinar cells, and then with hepatocytes, with whom they share endodermal origins. Conversely, endoderm-derived islet cells do not cluster with ectoderm-derived neurons^23^, despite common tissue-specific gene regulation and exocytosis machinery^24^.

Additional examples include the clustering of gastric, small intestine and colon epithelial cells; the clustering of all blood cell types; and the clustering of multiple mesoderm-derived cell types including vascular endothelial cells, adipocytes and skeletal muscle. Interestingly, lung bronchial epithelium clustered with esophagus and oral epithelium, whereas lung alveolar epithelium clustered with intestinal epithelium, consistent with evidence of early developmental origins of the alveolar cell lineage^25^.

Some methylation patterns were common to lineages that formed during early developmental stages. For example, 776 regions are unmethylated in epithelial cells derived from early endodermal derivatives, and methylated in mesoderm and ectoderm derived cells. We hypothesize these were demethylated in the endoderm germ layer, with derived cell types retaining these patterns decades later. Since endoderm derivatives do not share common function or gene expression, this provides yet another striking example of methylation patterns as a stable lineage mark.

Finally, we applied the same segmentation and clustering approach to a published methylation atlas from the Roadmap Epigenomics project^5^. The algorithm did not group related cell types, and often clustered samples based on donor identity. This further emphasizes the importance of careful purification of homogeneous cell types, avoiding mixed cell populations (Extended Fig. S5)

### Tens to hundreds of methylation blocks uniquely characterize each cell type

We next turned to study genomic regions that are differentially methylated in a cell type-specific manner. We organized the 207 samples into 39 groups of specific cell types, including blood cell types (B, T, NK, Granulocytes, monocytes and tissue-resident macrophages), breast epithelium (basal or luminal), lung epithelium (alveolar or bronchial), pancreatic endocrine (alpha, beta, delta) or exocrine (acinar and duct) cells, vascular endothelial cells from various sources, cardiomyocytes and cardiac fibroblasts, and more. We also defined 12 super-groups, where related cell types were grouped together, including muscle cells, gastrointestinal epithelium, pancreas, and more (Extended Table S3).

We then focused on differentially methylated blocks, composed of 5 CpGs or more, that are methylated (average methylation in block ≥66%) in one group of cell types, but unmethylated (≤33%) in all other samples, or vice versa. Overall, we identified 11,125 differentially methylated genomic regions. Intriguingly, almost all regions (98%, 10,892) were unmethylated in one cell type, and methylated in all others (see below).

To obtain a human cell type-specific methylation atlas, we identified the top 25 differentially unmethylated regions for each cell type (Figure 3 and Extended Table S4). These regions are uniquely demethylated in particular cell types and methylated in all other samples, and can serve as sensitive biomarkers for quantifying the presence of DNA from a specific cell type in a mixture. This approach has various applications, including the analysis of circulating cfDNA fragments^16–19,26–28^. Importantly, only <1% of the cell type-specific markers are covered by RRBS sequencing, 8-9% are covered by methyl-seq hybrid capture panels, and 13-22% are represented in the single-CpG 450K/EPIC arrays^11^, emphasizing the benefits of whole-genome sequencing for exhaustive identification of biomarkers.

**Figure 3.**
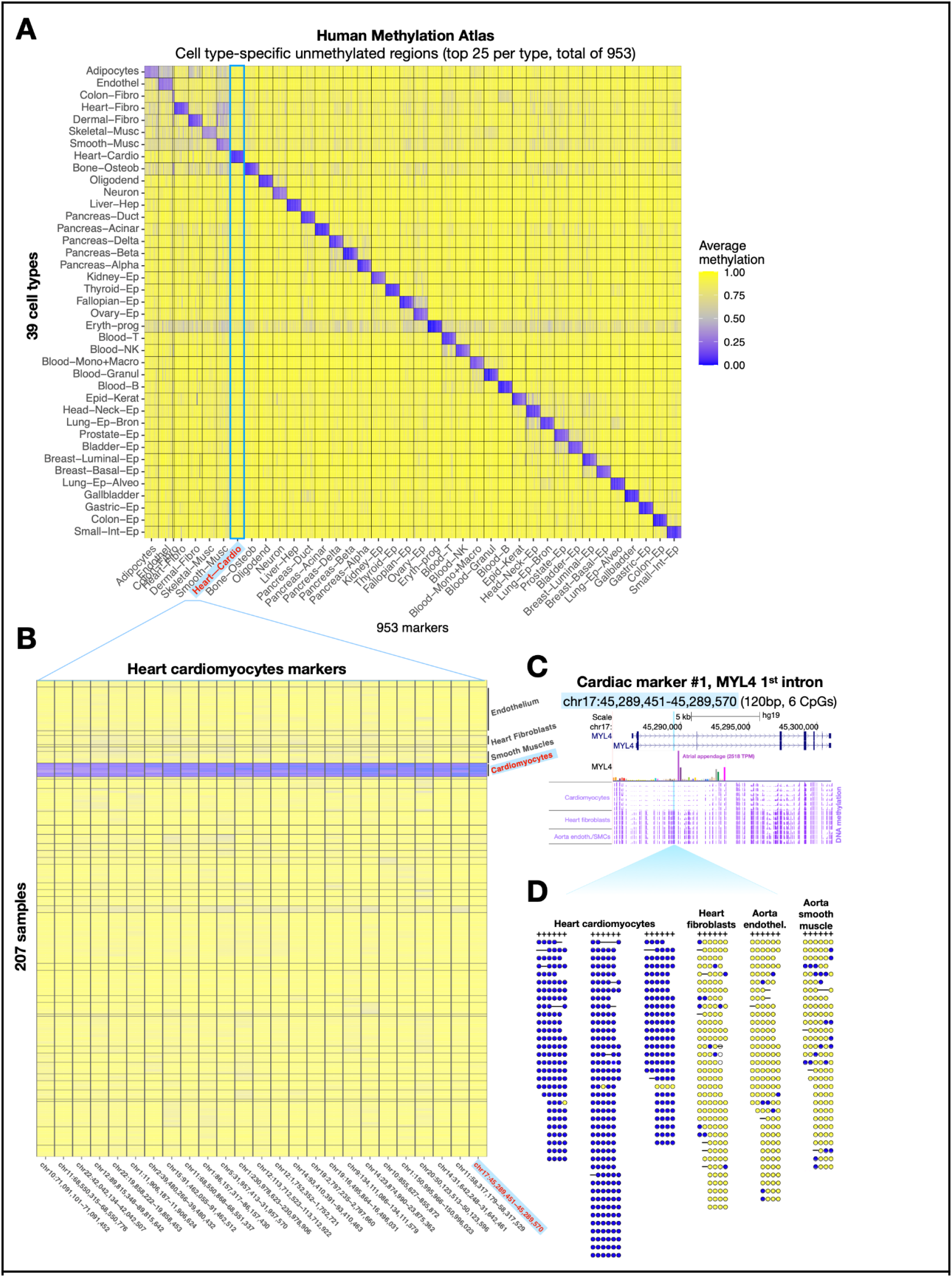
A Human Methylation Atlas of 207 samples across 39 cell types. **(A)** 953 genomic regions, unmethylated in a cell type-specific manner. Each cell in the plot marks the average methylation of one genomic region (column) at each of 39 cell types (rows). Up to 25 regions are shown per cell type, with a mean length of 251 bp (9 CpGs) per region. **(B) Top 25 cardiomyocyte regions**. For each region, we plot the average methylation of each CpG site (columns) across all 207 samples in the atlas, grouped into 39 cell types as before. **(C) A locus specifically unmethylated in cardiomyocytes**. This marker (highlighted in light blue) is 120bp long (6 CpGs), and is located in the first intron of MYL4, a heart-specific gene (TPM expression of 2518 in atrial appendage, GTEx inset). Genomic snapshot depicts average methylation (purple tracks) across six cardiomyocyte samples, four cardiac fibroblast samples, and three aorta samples (two endothelial, one smooth muscle cells). **(D) Visualization of bisulfite converted fragments** from three cardiomyocyte samples, one cardiac fibroblast sample, and two aorta samples (endothelium and smooth muscle). Shown are reads mapped to chr17:45289451-45289570 (hg19), with at least 3 covered CpGs. Yellow/blue dots depict methylated/unmethylated CpG sites.

### Cell type-specific unmethylated regions are tissue-specific enhancers

We next turned to characterize these sets of cell type-specific differentially unmethylated regions. Using GREAT, we identified the genes adjacent to each group of markers, and tested their enrichment for various gene-set annotations^29^. Genes adjacent to loci uniquely unmethylated in a given cell type typically reflected the functional identity of this cell type (Extended Table S5). For example, genes near B cell markers were enriched for B cell morphology, differentiation, IgM levels, and lymphopoiesis; NK cell markers were associated with NK cell mediated cytotoxicity, hematopoietic system, cytotoxicity, and lymphocyte physiology; Fallopian tube markers were enriched for egg coat and perivitelline space; and cardiomyocyte markers for cardiac relaxation, systolic pressure, muscle development, and hypertrophy.

We then analyzed the DNA accessibility and chromatin packaging of cell type-specific markers as defined by ATAC-seq, DNaseI-seq^5,30^ and histone marks indicative of active promoters and enhancers^5^. The top 250 unmethylated markers for monocytes and macrophages are highly accessible and characterized by H3K27ac and H3K4me1 in monocytes, whereas markers of other cell types show no enrichment in monocytes (Figure 4A), with similar results for markers of other cell types (Extended Figure S6). We also show strong coordinated enrichment of chromHMM enhancer annotations at cell type-specific markers31 (Figure 4). These findings are consistent with previous studies that have associated tissue-specific demethylation with gene enhancers^1,32^.

**Figure 4.**
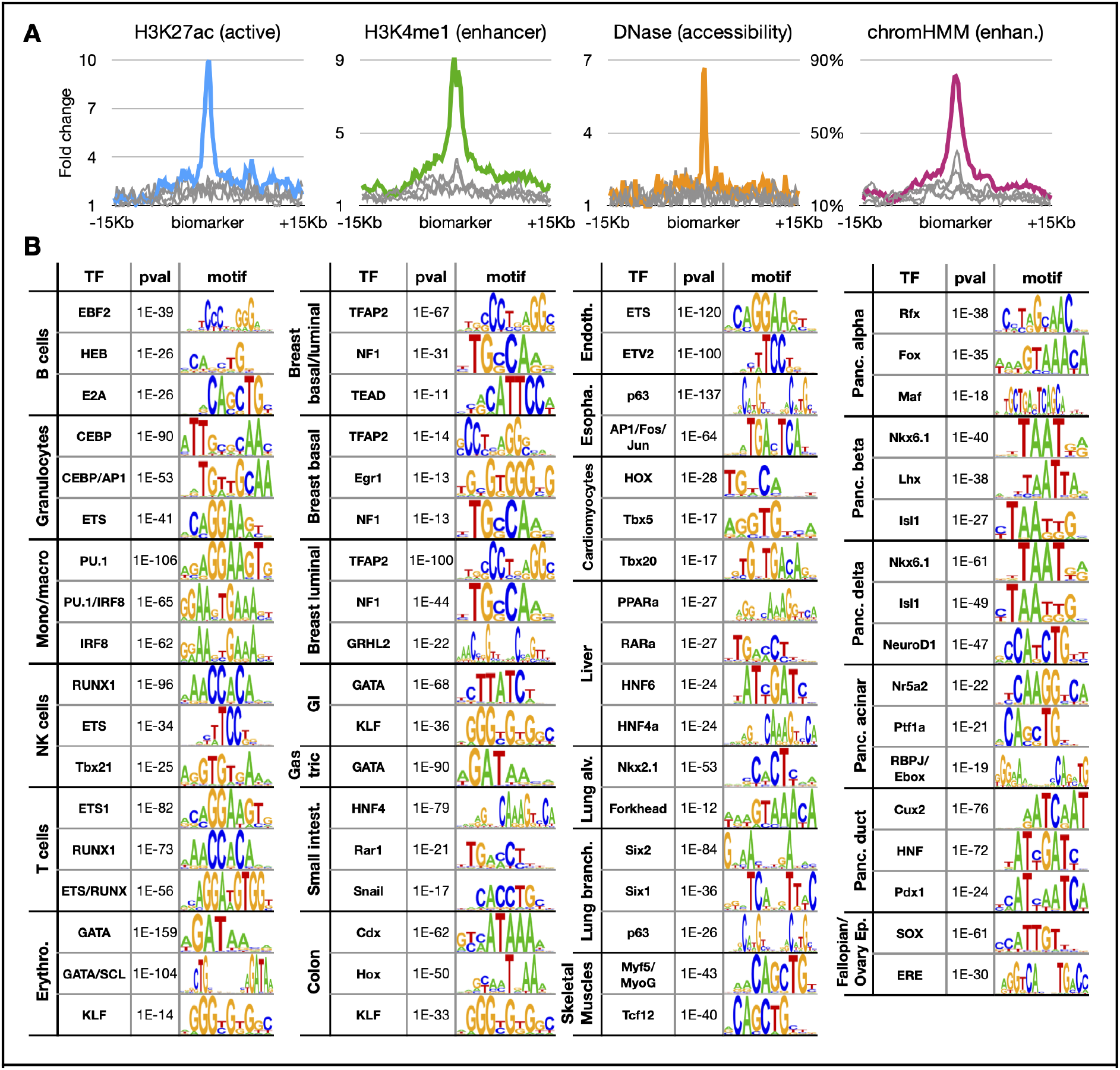
Cell type-specific markers as putative enhancers. **A.** Average ChIP-seq signal for the active regulatory mark H3K27ac, enhancer mark H3K4me1, DNA accessibility, and chromHMM enhancer annotations for the top 250 cell-type specific unmethylated markers for Monocyte/Macrophages. The average signal for top 250 markers of other blood cell types (Granulocytes, B, T, and NK cells) are shown as grey lines, for comparison. **B.** Cell type-specific markers are enriched for regulatory motifs. Shown are the top transcription factor binding site motifs, enriched among the top 1,000 differentially unmethylated regions per cell type, using HOMER motif analysis. Motifs similar to prior (more significant) hits are skipped.

To further assess the biological importance of cell type-specific unmethylated regions, we studied their association with transcription factors (TF) that could affect DNA methylation, or bind DNA in a cell type-specific manner, depending on methylation and chromatin^33–36^. We performed motif analysis using HOMER^37^, and calculated the enrichment (within the unmethylated markers of each cell type) for known TF binding motifs (Extended Table S6). For most cell types the top motifs included master regulators and key TF (Figure 4B). For example, B cells are enriched for Ebf2/HEB/E2A, granulocytes for CEBP/AP1/ETS, and T cells for ETS/RUNX. This association between cell type-specific unmethylated regions and TF binding motifs could identify novel gene regulatory circuits and expose distal enhancers active in specific cell types.

### Identification of target genes regulated by cell-type specific enhancers

We aimed to identify the target genes of putative enhancers marked by cell type-specific demethylation. Often, top markers fall within intronic regions and are likely to regulate these genes (for example glucagon in pancreatic alpha-cells; NPPA, MYH6, and MYL4 in cardiomyocytes, or EPCAM in GI epithelial cells), or proximally to likely targets (e.g., a beta-cell marker 5Kb from the Insulin gene). Other markers are further apart from their target genes. We devised a computational algorithm that integrates the distance between a marker and its surrounding genes, with their expression patterns across multiple cell types. We applied an iterative bidirectional z-score calculation, where the expression level of a gene in a given cell type is compared to the expression of other genes, and the expression of that gene elsewhere. This highlighted hallmark genes for many cell types, and suggested putative targets for many of the top 25 unmethylated markers for each cell type. For example, hepatocyte markers were associated with APOE, APOC1, APOC2, Alpha 2-antiplasmin, and glucagon receptor (GCGR). Similarly, cardiomyocyte markers were associated with NPPA, NPPB, and myosin genes; and pancreatic islet markers with the insulin and glucagon genes (Extended Table S7). These findings further support the principle that loci specifically unmethylated in a given cell type are likely enhancers positively regulating genes expressed in this cell type, often controlling adjacent genes. We note however genes adjacent to a locus specifically unmethylated in a given cell type are often broadly expressed beyond this cell type (see discussion).

### Cell type-specific hyper-methylated regions are enriched for CpG islands and for polycomb, CTCF, and REST targets

We studied the genomic regions that are methylated in one cell type, but unmethylated elsewhere in the human body. These are enriched for CpG islands (38% of methylated regions, compared to 1.7-2.7% of cell type-specific unmethylated regions), and are marked by H3K27me3 and polycomb in other cell types (Figure 5A-C), as previously reported for cancer and developmental processes^38,39^. These cell type-specific hyper-methylated regions were generally less significant for motif enrichment (compared to uniquely unmethylated regions). Intriguingly, only ~3% of the total set of cell type-specific differentially methylated regions are hyper-methylated.

**Figure 5.**
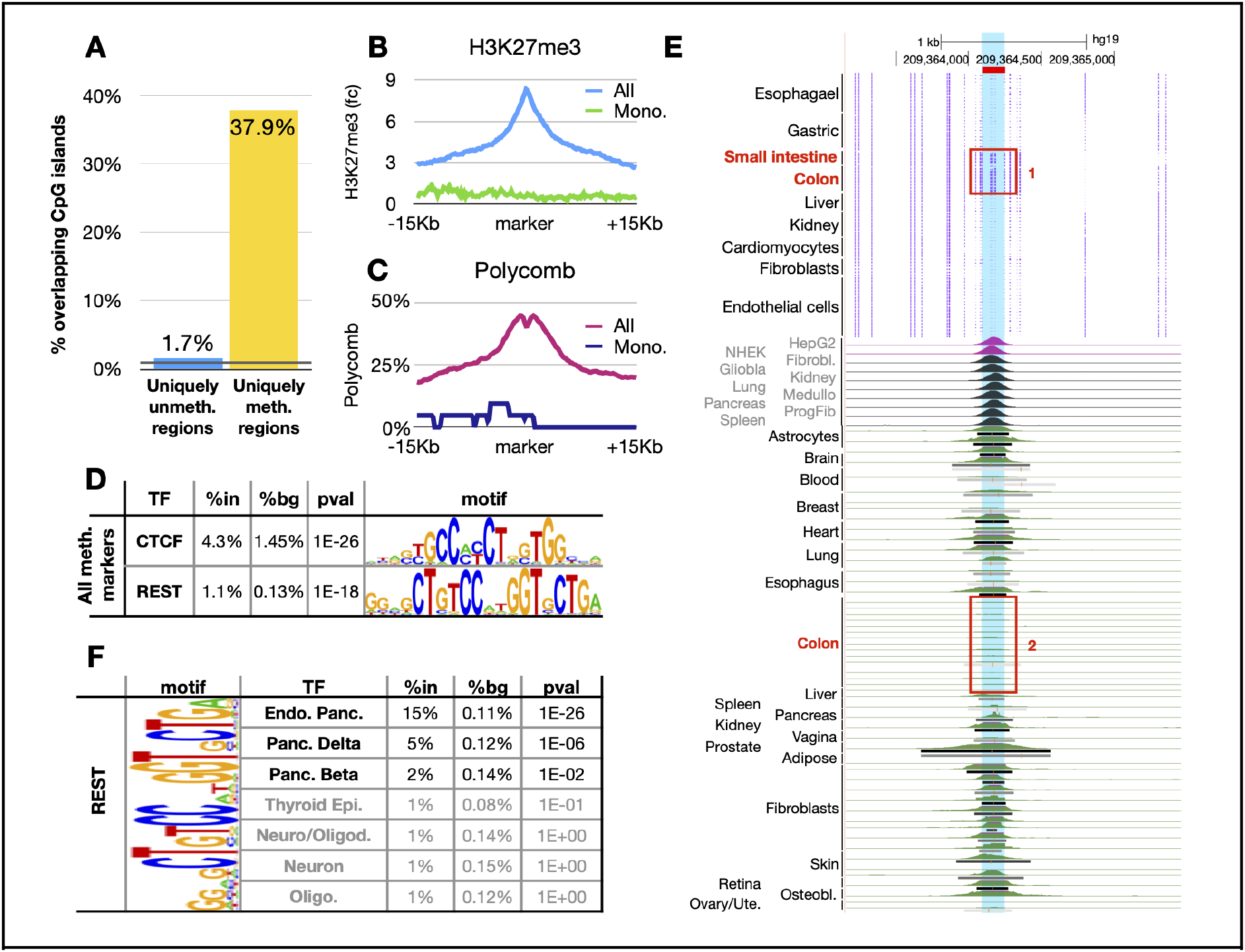
Cell type-specific hyper-methylated regions are enriched for CpG islands, polycomb targets, and CTCF and REST/NSRF. **(A)** 37.9% of top cell type-specific hyper-methylated markers (1,185 of 3,125, p<1E-100) overap CpG islands. For comparison, 1.7% of cell type-specific hypo-methylated regions (198 of 11,371, p<2E-29) overlap CpG islands, which make <0.9% of the genome (black line). **(B)** These regions are typically enriched for H3K27me3 in other cell types. Shown are the average H3K27me3 signals in monocytes and macrophages near all cell type-specific hyper-methylated regions (top, blue) or near monocytes/macrophages-specific hyper-methylated regions (green). **(C)** Similar plots for Polycomb annotations in monocytes and macrophages (chromHMM), for all or monocyte/macrophage-specific markers. **(D)** Motif analysis of cell type-specific hyper-methylated regions (top 100 per cell type) identifies known CTCF and REST/NSRF motifs. **(E)** Analysis of ChIP-seq data for one such site (chr1:209364093-209364250, highlighted in blue, hg19), specifically methylated in the small intestine and colon epithelium (box 1), and unmethylated elsewhere. As shown below, this site is bound in multiple cell types and tissues, but is mostly unbound in the stomach and colon epithelium, in vivo (box 2). **(F)** REST/NSRF motif is present within 15% of top 100 cell type-specific hyper-methylated regions in the endocrine pancreas, 5% of top delta-cell markers, and 2% of top beta-cell markers, compared to ~0.1% in background sequences, in accordance with REST target expression in the endocrine pancreas.

After pooling together all cell type-specific hyper-methylated regions, we identified a strong enrichment for target sequences of the chromatin regulator CTCF (p≤1E-26, Figure 5D). This suggests that DNA methylation of CTCF binding sites could act as a tissue-specific regulatory switch to modulate its binding, potentially affecting tissue-specific 3D genomic organization^33,40,41^. To test this idea, we compared patterns of DNA methylation at CTCF sites with genome-wide CTCF protein binding in specific tissues. Figure 5E shows the methylation pattern and the published *in vivo* CTCF occupancy at one locus, which is methylated specifically in the colon and intestine. Consistent with DNA methylation preventing CTCF binding, ChIP data show selective absence of CTCF binding at this locus in the colon. In addition, loci methylated in specific cell types were enriched for targets of the transcriptional repressor of neural genes, REST/NRSF (p≤1E-18), and this was seen most prominently in the methylome of pancreatic islet cells (Figure 5F). While DNA methylation has not been shown to affect the binding or activity of REST, this finding raises the intriguing possibility that methylation of REST targets in islets could permit endocrine differentiation independently of REST repression.

### Accurate and sensitive cellular decomposition outperforms array-based atlas

Lastly, we applied the methylation atlas for analysis of methylomes obtained from composite tissue samples and cfDNA. We developed a computational framework for labeling sequenced DNA fragments based on their methylation status, computed the methylation score of each biomarker across all cell types, and applied a computational deconvolution algorithm for the analysis of methylomes derived from mixtures.

To estimate the accuracy of our markers, we used *in silico* mixtures of sequenced reads. For each cell type, we mixed a given percentage of reads from that cell type in leukocyte reads, then used the deconvolution algorithm to infer the amount of each cell type in the mixture. We repeated this process for all cell types at concentrations varying from 0% to 10%. As shown in Figure 6A, we found that a set of 25 methylation markers per cell type resulted in an ability to accurately detect DNA from a given source when present in just 0.03-0.1% of a mixture, an order-of-magnitude improvement in comparison to current 450K approaches.

**Figure 6.**
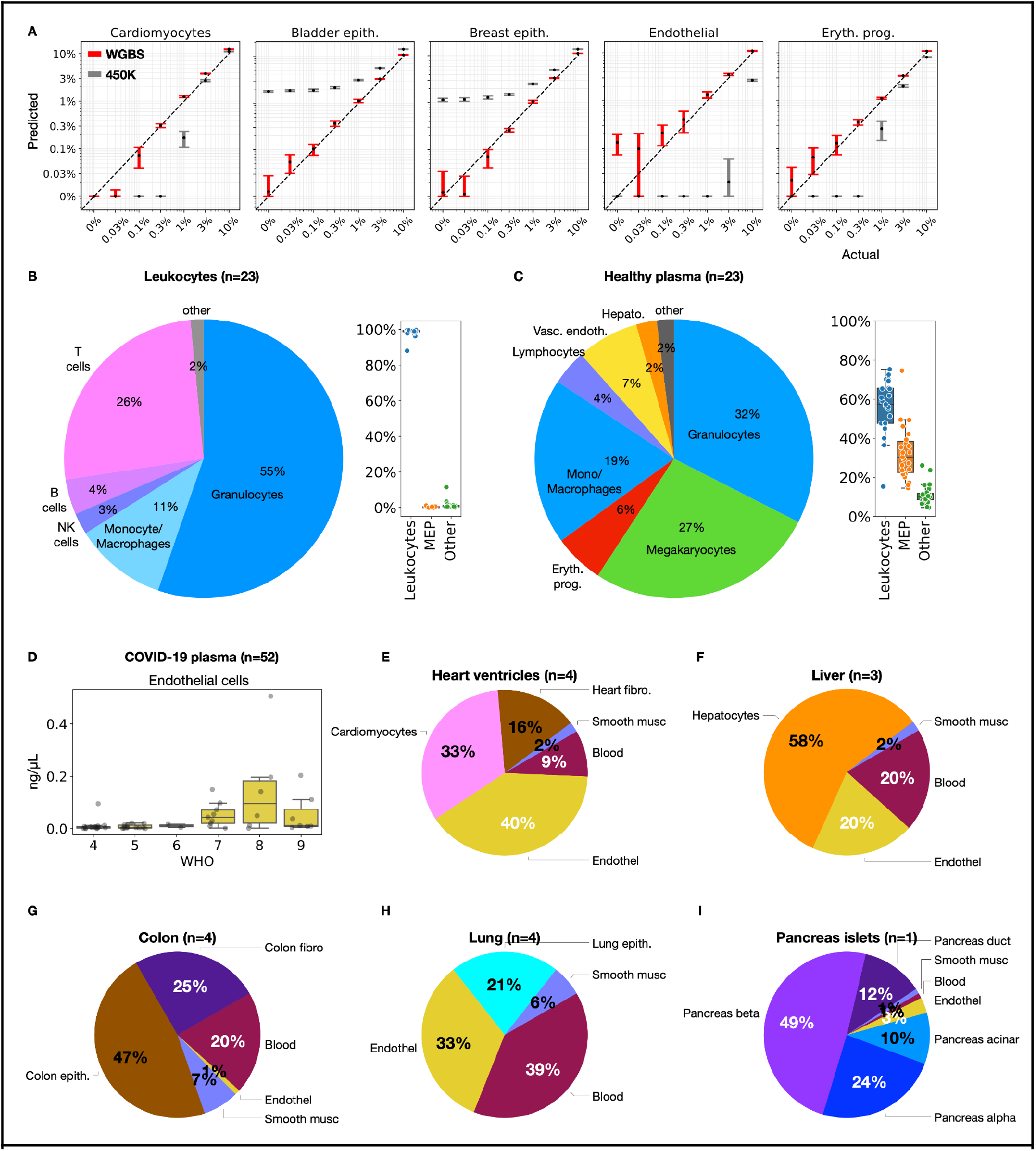
Fragment-level deconvolution using cell type-specific biomarkers. **(A)** Cell type-specific markers outperform array-based atlas and achieve <0.1% resolution. Shown are *in silico* simulations for five cell types, computationally mixed within leukocytes. Each mixture was analyzed using our atlas (red), and compared to Moss et al. (gray). Box plots show predicted contribution in 10 simulations, with 1 SD error bars. **(B-C)** Cell type composition in leukocytes and plasma samples from healthy donors. Box plots show overall proportions of leukocytes, megakaryocytes and erythroblasts (MEP), and other cell types. **(D)** Analysis of low coverage plasma samples from 52 SARS-CoV-2 patients^42^ identifies endothelial derived cfDNA in patients with WHO ordinal scale ≥ 7 (requiring admittance to ICU). **(E-I) Fragment-level deconvolution of Roadmap/ENCODE samples**^5,6^ **reveals cell type-specific contributions** (E) Heart ventricle samples contain a mixture of cardiomyocytes, endothelial cells, fibroblasts, and blood. (F) Liver samples contain <60% of hepatocyte DNA, with blood and endothelial cells. (G) Colon samples are composed of <50% epithelium, with fibroblasts and blood. (H) Lung sample contains <30% of lung epithelial cells. (I) Pancreas islets show composition of beta, alpha, duct and acinar cells.

We then estimated the cellular composition of leukocytes and cfDNA using WGBS data from 23 healthy donors. Over 98% of leukocyte-derived DNA was attributed to granulocytes, monocytes and macrophages, NK, T, and B cells, consistent with typical blood counts (Figure 6B, Extended Table S8). The cfDNA of healthy subjects was mostly derived from leukocytes: granulocytes (32%), monocytes/macrophages (19%), and lymphocytes (4%). Solid tissues that contributed to cfDNA included vascular endothelial cells (7%) and hepatocytes (2.5%) (Figure 6C), consistent with previous results^26^. The current atlas also revealed a significant contribution of megakaryocytes (27%) and erythrocyte progenitor cells (6%) to cfDNA, which was not observed in previous studies that used reference methylomes of a more limited scope.

### Endothelial cell-free DNA correlates with COVID-19 severity

Analysis based on DNA methylation patterns offers an opportunity to identify the tissue origins of cfDNA. COVID-19 inflicts damage to multiple tissues, some of which have no biomarkers. We used the atlas to deconvolve shallow whole-genome bisulfite sequencing data from 52 hospitalized COVID-19 patients^42^. We identified excessive cell-free DNA fragments from granulocytes, erythrocyte progenitors, lung and liver, consistent with published analysis of these samples. Strikingly, we also identified a significant contribution of vascular endothelial cells to the cfDNA of these patients, which could not be discovered in the published analysis in the absence of endothelial cell methylome reference (Figure 6D). Interestingly, the concentration of endothelial cell-derived cfDNA was higher in patients with a severe disease (WHO score≥7), compared to those with a milder disease (WHO score≤6, p≤6E-5, Mann-Whitney). These results suggest that vascular endothelial cell death plays a significant role in the pathogenesis of COVID-19, potentially related to coagulopathy, and highlight the benefit of using a comprehensive cell type-specific atlas for cfDNA methylome analysis.

### Cell-type disentanglement of composite samples

Finally, we analyzed whole-genome methylomes from ENCODE^6^ and the Roadmap Epigenomics atlas^5^ using our atlas (based on 25 markers per cell type). Deconvolution of some methylomes revealed a homogenous composition as intended, e.g. 99% T-cell DNA in Roadmap T cells samples (Extended Table S9). However analysis of other samples revealed a highly heterogeneous composition. For example, heart ventricle samples were composed of only 33% cardiomyocytes, 40% endothelial cells and 16% cardiac fibroblasts (Fig. 6E); one pancreatic islet methylome had as much as 22% exocrine pancreas; liver methylomes were made of <60% hepatocytes, 20% blood and 20% endothelial cells; and the colon methylomes with <50% colon epithelium, 25% colon fibroblasts and 20% blood. Most strikingly, Roadmap lung samples are dominated by blood (39%), endothelium (33%), and smooth muscle (6%), with only 7-29% of DNA derived from lung epithelial cells (Figure 6F-I and Extended Table S9). Thus, the atlas facilitates determination of tissue composition, an essential requirement for extraction of accurate information about methylation patterns in any given sample.

## Discussion

The comprehensive atlas of human cell type methylomes described here sheds light on principles of DNA methylation, and provides a valuable resource for multiple lines of investigation, as well as translational applications.

### Variation of DNA methylation between replicates and different cell types

Our analysis revealed that methylation patterns are strikingly similar among healthy replicates of the same cell type from different individuals. The similarity between individuals reflects the extreme robustness of cell differentiation and maintenance circuits, at least as far as healthy tissues are concerned. Pathologies involving destabilization of the epigenome obviously disrupt these circuits, resulting in a larger variety of methylation patterns among cells that descend from a specific normal cell type. We predict that even in cancers (of the same primary anatomic site and histologic type), comparative methylome analysis of purified epithelial cells, performed at the level of methylation blocks, will reveal a smaller inter-individual variation than typically assumed.

As the atlas blocks revealed, each cell type has a set of genomic regions that are uniquely unmethylated in that cell type compared to others, as well as additional genomic regions that share methylation patterns with related cell types. Using unsupervised clustering of cell type-specific methylomes we found that cell types in the atlas were clustered in ways that reflected their developmental origins, rather than expression patterns. This offers a fascinating view of DNA methylation as a living record of the methylomes of progenitor cells, retained in the genome through dramatic developmental transitions and decades of life thereafter. Perhaps the most striking example of this principle is the clustering of cells according to their germ layer of origin. The loci that drive the clustering of colon epithelial cells from one adult donor with lung alveolar cells of another donor are probably reflecting the common origins of these cell types in the embryonic endoderm, which forms during gastrulation and diverges shortly after. We propose that comparative methylome analysis will allow reconstructing parts of the methylomes of fetal structures or cell types, similarly to the reconstruction of last common ancestors in evolutionary biology.

### Cell type-specific demethylation identifies enhancers and TF binding motifs

The vast majority of the cell type-specific differentially methylated regions were specifically demethylated in one cell type. The chromatin of these regions is typically highly accessible and bears histone marks associated with active gene regulation, as found in enhancers and promoters. Moreover, differentially unmethylated loci are enriched for TF binding site motifs that operate in that cell type. Finally, we devised an integrated approach that, based on distance and gene expression profiles, allowed us to highlight possible target genes for these putative enhancer regions. Notably, many enhancer regions were associated with nearby genes that are broadly expressed, potentially reflecting gene regulation by multiple tissue-specific enhancers. Our findings are consistent with, and considerably expand upon, previous studies that revealed tissue-specific hypomethylation occuring at enhancers^33–35^. Further analysis of this atlas will reveal the complete set of human enhancers in each cell type.

### Roles for cell type-specific hyper-methylation

Conversely, we identified genomic regions that are specifically methylated in one or two cell types, representing ~3% of cell type-specific differentially methylated regions. They are often located in CpG islands, and characterized by H3K27me3 and polycomb binding in tissues where the locus is not methylated^38,39^. This epigenetic repressive switching was previously described in cancer and during early development^39,43^, but its role during differentiation of specific cell types remains unclear. These regions are enriched for CTCF binding sites, suggesting a role for DNA methylation in attenuating the binding of CTCF and thus modulating the cell type-specific 3D organization of neighboring DNA^33,34,44^.

### Cell type-specific DNA methylation biomarkers for cell-free DNA analysis

The atlas described here is the most comprehensive whole-genome healthy DNA methylation atlas to-date. We have identified over a thousand cell type-unique DNA methylation regions that could serve as accurate and specific biomarkers for identifying cell death events by monitoring cfDNA. Notably, most of these marker regions are not covered by 450K/EPIC BeadChip DNA methylation arrays, and were not previously appreciated. As may be apparent, many cell types are missing in the atlas, typically because of limited availability of material. Examples include osteoblasts, cholangiocytes, cells of the adrenal gland, urethral epithelium and hematopoietic stem cells. Additionally, we did not separate many sub-populations of interest, for example different types of neurons or lymphocytes. The atlas is viewed as a living, publicly available database to be updated in the future. The resolution of the atlas yields a quantitative understanding of composite tissues, and allows one to identify missing methylomes of additional cell types that are yet to be characterized.

In summary, we present a comprehensive methylome atlas of primary human cell types and provide examples for biological insights that can be gleaned from this resource. Among the many potential utilities of this atlas, perhaps most promising is the possibility to use it for deconvolution of cell types in a mixed cell type sample, and sensitive identification of the tissue of origin of cfDNA in plasma of individuals with cancer and other diseases^16–19,26–28^.

## Methods

### Human tissue samples

Human tissues were obtained from various sources as detailed in Extended Table S1. The majority (150) of the 207 samples analyzed were sorted from tissue remnants obtained at the time of routine, clinically indicated surgical procedures at the Hadassah Medical Center (Extended Table S1). In all cases, normal tissue distant from any known pathology was used. Surgeons and/or pathologists were consulted prior to removing the tissue to confirm that its removal would not compromise the final pathologic diagnosis in any way. For example, in patients undergoing right colectomy for carcinoma in the cecum, the distal most part of the ascending colon and the most proximal part of the terminal ileum were obtained for cell isolation. Normal bone marrow was obtained at the time of joint replacement in patients with no known hematologic pathology. The patient population included 137 individuals (n=61 males, n=75 females), aged 3-83 years. The majority of donors were White. Approval for collection of normal tissue remnants was provided by the Institutional Review Board (IRB, Helsinki Committee), Hadassah Medical Center, Jerusalem, Israel. Written informed consent was obtained from each donor or legal guardian prior to surgery.

As described in Extended Table S1, some cells and tissues were obtained through collaborative arrangements: pancreatic exocrine and liver samples (cadaveric organ donors, n=5) from Prof. Markus Grompe, Oregon Health & Science University. Adipocytes (subcutaneous adipocytes at time of cosmetic surgery following weight loss; n=3), oligodendrocytes and neurons (brain autopsies, n=14) from Profs. Kirsty L. Spalding and Henrik Druid, Karolinska Institute, Stockholm, and research grade cadaveric pancreatic islets from Prof. James Shapiro, University of Alberta (n=16). In all cases tissues were obtained and transferred in compliance with local laws and after the approval of the local ethics committee on human experimentation. Sixteen cell types were obtained from commercial sources, including 15 from Lonza Walkersville, Walkersville, MD, U.S.A. and one from Sigma Aldrich. Three cell preparations were obtained from the Integrated Islet Distribution Program (IIDP, https://iidp.coh.org).

### Tissue dissociation and FACS sorting of purified cell populations

Fresh tissue obtained at the time of surgery was trimmed to remove extraneous tissue. Cells were dispersed using enzyme-based protocols optimized for each tissue type (Extended Figure S1). The resulting single-cell suspension was incubated with the relevant antibodies and FACS sorted to obtain the desired cell type.

Purity of live sorted cells was determined by mRNA analysis for key known cell-type specific genes whereas purity of cells that were fixed prior to sorting was determined using previously validated cell-type specific methylation signals. DNA was extracted using the DNeasy Blood and Tissue kit (#69504 Qiagen; Germantown, MD) according to the manufacturer’s instructions, and stored at −20°C for bisulfite conversion and whole genome sequencing.

### Whole-genome bisulfite sequencing

Up to 75 ng of sheared gDNA was subjected to bisulfite conversion using the EZ-96 DNA Methylation Kit (Zymo Research; Irvine, CA), with liquid handling on a Hamilton MicroLab STAR (Hamilton; Reno, NV). Dual indexed sequencing libraries were prepared using Accel-NGS Methyl-Seq DNA library preparation kits (Swift BioSciences; Ann Arbor, MI) and custom liquid handling scripts executed on the Hamilton MicroLab STAR. Libraries were quantified using KAPA Library Quantification Kits for Illumina Platforms (Kapa Biosystems; Wilmington, MA). Four uniquely dual indexed libraries, along with 10% PhiX v3 library (Illumina; San Diego, CA), were pooled and clustered on a Illumina NovaSeq 6000 S2 flow cell followed by 150-bp paired-end sequencing.

### Whole-genome bisulfite sequencing computational processing

Paired-end FASTQ files were mapped to the human (hg19), lambda, pUC19 and viral genomes using bwa-meth (V 0.2.0), with default parameters^45^, then converted to BAM files using SAMtools (V 1.9)^46^. Duplicated reads were marked by Sambamba (V 0.6.5), with parameters “-l 1 -t 16 --sort-buffer-size 16000 --overflow-list-size 10000000”^47^. Reads with low mapping quality, duplicated, or not mapped in a proper pair were excluded using SAMtools view with parameters -F 1796 -q 10. Reads were stripped from non-CpG nucleotides and converted to PAT files using *wgbstools* (V 0.1.0)^48^.

### Genomic segmentation into multi-sample homogenous blocks

We developed and implemented a multi-channel Dynamic Programming segmentation algorithm to divide the genome into continuous genomic regions (blocks) showing homogeneous methylation levels across multiple CpGs, for each sample^48^. We model the CpG sites with a generative probabilistic model, assuming there is a universal underlying segmentation of all ~28M sites into an unknown number of blocks. This segmentation, unlike the methylation patterns, is similar across different cell types and individuals. Each block induces a Bernoulli distribution with some θ_*i*_^*k*^, where *i* is the block index and *k* is the sample (*k* = 1,…, *K*), and all CpG sites are represented by random variables sampled i.i.d from the same beta value *Ber*(θ_*i*_^*k*^).

### Finding an Optimal Segmentation

We used dynamic programming to find a segmentation that maximizes a log-likelihood score for the blocks. The score for the i’th block is the log-likelihood of the beta values of the sites in this block across all K samples. We compute K Bayesian estimators for the block’s parameters 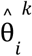:

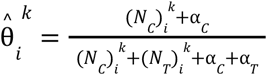

Where (*N_C_*)_*i*_^*k*^, (*N_T_*)_*i*_^*k*^ is the number of observations of sites in the block i and sample k that are methylated/unmethylated. α_*C*_, α_*T*_ are pseudocounts for methylated/unmethylated observations in block i. They are constant hyper-parameters of the model, which set the tradeoff between longer to more homogenous blocks. The log-likelihood of a single block in a single example is:

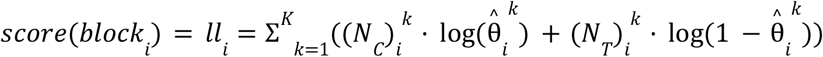

### Dynamic Programming Algorithm

We maintain a 1 × *N* table T for the optimal scores (N=28,217,448). *T*[*i*] holds the score of the optimal segmentation of sites 1,…, *i*. *T*[*N*] holds the final optimal score. The table is updated from 1 to N as follows:

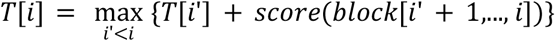

*T*[*i*] is the maximum over the sites preceding site *i*(*i*’ < *i*), of the score of the optimal segmentation that ends on site *i*’(*T*[*i*’]), concatenated with the single block from *i*’ + 1 to *i*. A similar traceback table is also maintained, in order to retrieve the optimal segmentation. In order to improve performance, we set an upper bound on block length (either in CpG sites or bases), which improves running time and allows for multi-processing.

### Segmentation and clustering analysis

We segmented the genome into 7,264,350 blocks using wgbstools (with parameters “segment --max_bp 5000”), with all of the 207 samples as reference, and retained 2,107,635 blocks that cover ≥4 CpGs. For the hierarchical clustering we selected the top 1% (21,077) blocks showing the largest variability in average methylation across all samples. Blocks with sufficient coverage of ≥10 observations (calculated as sequenced CpG sites) across ⅔ of the samples we further retained. We then computed the average methylation for each block and sample calculated using wgbstools (“--beta_to_table -c 10”), marked blocks with <10 observations as missing values, and imputed their methylation values using sklearn KNNImputer (V 0.24.2)^49^. The 207 samples were clustered with the UPGMA clustering algorithm^21^, using scipy (V 1.6.3)^50^, and L1 norm. The fanning diagram (Figure 2) was plotted using ggtree (V 2.2.4)^51^.

### Cell type-specific markers

The 207 atlas samples were grouped into 51 groups by their cell type (Extended Table S3), including 39 basic groups (e.g. epithelial cells pancreatic alpha-cells), and composite super-groups (e.g. epithelial alpha, beta, and delta cells, all from the endocrine pancreas). We performed a one-vs-all comparison, to identify differentially methylated blocks unique for each set. For this, we first identified blocks that cover ≥5 CpGs, with length varying between 10 to 500bp. We then calculated the average methylation per block/sample, as the number of methylated CpGs sites within all sequenced reads across each block). Blocks with insufficient coverage (<25 observations) were assigned a default value of 0.5. We then selected blocks with average methylation below 0.33 across samples from one cell type, with an average methylation of ≥0.66 in all others, or vice versa.

For cell type-specific markers, we selected the top 25, 100 or 250 or 1,000 blocks with the highest delta-beta for each cell type. For hypo-methylated markers this was computed as the difference between the 75^th^ percentile among the block average methylation within samples in the target set, and the 2.5^th^ percentile among the rest of samples (background set). This allowed for ~1 outlier sample in the target group, and ~5 outliers outside. Analogously, for hyper-methylated markers we computed the 97.5^th^ percentile of the background and the 25^th^ percentile within the target samples.

### Enrichment for gene set annotations

Analysis of gene set enrichment was performed using GREAT^29^. For each cell type, we selected the top 250 differentially unmethylated regions, and ran GREAT via batch web interface using default parameters. Enrichments for “Ensembl Genes” were ignored, and a significance threshold of Binomial FDR≤0.05 was used.

### Enrichment for chromatin marks

For each cell type, we analyzed the top 250 differentially unmethylated regions vs. published ChIP-seq (H3K27ac and H3K4me1) and DNase-seq from the Roadmap Epigenomics project (downloaded from ftp.ncbi.nlm.nih.gov/pub/geo/DATA/roadmapepigenomics/by_experiment and http://egg2.wustl.edu/roadmap/data/byDataType/dnase/BED_files_enh in bigwig and bed formats). These include E032 for B cells markers, E034 for T cells markers, E029 for monocytes/macrophages markers, E066 for liver hepatocytes, E104 for heart cardiomyocytes and fibroblasts, E109 and E110 for gastric/small intestine/colon^5^. Annotations for chromHMM were downloaded (15-states version) from https://egg2.wustl.edu/roadmap/data/byFileType/chromhmmSegmentations/ChmmModels/coreMarks/jointModel/final^31^, and genomic regions annotated as enhancers (7_Enh) were extracted and reformatted in bigwig format. Raw single-cell ATAC-seq data were downloaded from GEO GSE165659^30^, as “feature” and “matrix” files for 70 samples. For each sample, cells of the same type were pooled together to output a bedGraph file, which was mapped from hg38 to hg19 using UCSC liftOver^52^. Overlapping regions were dropped using bedtools (V 2.26.0)^53^. Finally, bigWig files were created using bedGraphToBigWig (V 4)^54^. Heatmaps and average plots were prepared using deepTools (V 3.4.1)^55^, with the computeMatrix, plotHeatmap, and plotProfile functions. We used default parameters except for --referencePoint=center, 15Kb margins, and binSize=200 for ChIP-seq, DNaseI and chromHMM data, and 75Kb margins with binSize=1000 for ATAC-seq data.

### Motif analysis

For each cell type, we analyzed the top 250 differentially unmethylated regions for known motifs, using HOMER’s findMotifsGenome.pl function, with -bits and -size 250 parameters^37^. Additionally, we analyzed the top 100 differentially methylated regions for each cell type (using the same parameters), as well as their combined set, composed of 3,125 regions in total.

### Methylation marker-gene associations

For each cell type-specific marker, we identified all neighboring genes, up to 500Kb apart. We then examined the expression levels of these genes across the GTEx datasets, covering 50 tissues and cell types^56^. We then calculated the over-expression level of each <gene, condition> pair, by computing the deviation (Z-score) of that gene across all conditions (row-wise calculation), and then the deviation of that condition across all genes (column-wise calculation), repeatedly until convergence. This Z-score reflects the bidirectional enrichment of each <gene, condition> combination, compared to all other genes/conditions. We then classified each <marker, gene, condition> combination as Tier 1: distance≤5Kb, expression≥10 TPM, and Z-score≥1.5; or Tier 2: same but as Tier 1, with dist≤50Kb; or Tier 3: up to 750Kb, expression≥25 TPM, and Z-score≥5; or Tier 4: same as Tier 3 with Z-score≥3.5.

### Inter-individual variation in cell type methylation

We define a similarity score between two samples as the fraction of blocks containing ≥3 CpGs, and ≥10 binary observations (sequenced CpG sites), where the average methylation of the two samples differs by ≥0.5. Only cell types with n≥3 FACS-sorted replicates from different donors are considered (138 samples in total).

### CTCF ChIP-seq analysis

CTCF ChIP-seq data were downloaded from the ENCODE project^6^, as 168 bigwig files, covering 61 tissues/cell types (hg19). Samples of the same cell type were averaged using multiBigwigSummary bins (V 3.4.1)^55^.

### Endodermal marker analysis

The 776 endodermal hypo-methylated markers were found using wgbstools’ find_markers function (V 0.1.0), with parameters “--delta 0.4 --tg_quant 0.1 --bg_quant 0.1”^48^. Endoderm-derived epithelium (51 samples was compared to 105 non-epithelial samples from mesoderm or ectoderm. Blocks were selected as markers if the average methylation of the 90^th^ percentile of the epithelial samples was lower than the 10^th^ percentile of the non-epithelial samples by at least 0.4.

### UXM fragment-level deconvolution algorithm

We developed a fragment-level deconvolution algorithm: the fraction of methylated CpG sites was computed per fragment, and it was annotated as U (mostly unmethylated), M (mostly methylated) or X (mixed). We then calculated, for each genomic region (marker), the proportion of U/X/M reads among all reads with ≥ k CpGs. Here, we used k=4 and thresholds of ≤25% methylated CpGs for U reads and ≥75% methylated CpGs for M. We then constructed a reference atlas *A* with 1,291 markers (top 25 markers per cell type/group, including megakaryocytes), wherein the *A_i,j_* cell holds the U proportion of the i’th marker in the j’th cell type (for hypo-methylated markers), or the M proportion (for hyper-methylated markers). Given an input sample, we compute the U or M read count within each marker as a 1,266×1 vector, we normalized each row in *A* accordingly, by multiplying the U/M proportions in each cell type by the total read coverage *C_i_* of each marker *i* in the input sample. Finally, non-negative least squares (NNLS) is applied to infer the coefficient vector *x* by minimizing |*diag*(*C*) · *A* · *x* − *b*|_2_ subject to non-negative x, and *x* is normalized such that 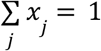.

### in silico simulations WGBS deconvolution

For each of the 39 cell types from the atlas, fragments were merged across replicates (after deduping) and split into training (70%) and test (30%) sets. For background, we merged n=23 leukocyte samples and simulated mixtures of different cell types, including cardiomyocytes (n=6), bladder epithelium (n=5), breast epithelium (n=7), endothelial cells (n=19), and erythrocyte progenitors (n=3). For each cell type, we simulated 10 mixtures at input proportions of 10%, 3%, 1%, 0.3%, 0.1%, 0.03% and 0%. Merging, splitting, and mixing of reads were performed using wgbstools [V0.1.0]^48^. A matching reference atlas was constructed using training set fragments only, across top 25 markers for each cell type. Simulated mixtures were analyzed using our UXM algorithm, across fragments with ≥5 CpGs in breast epithelial samples, and ≥4 for the other types.

Array based 450K data were simulated using wgbstools (beta_to_450k function, V0.1.0), and deconvolution performed as in Moss et al.26 (https://github.com/nloyfer/meth_atlas).

### WGBS deconvolution

Leukocytes and matching plasma samples (n=23) were processed as described above, and analyzed using the WGBS methylation atlas, including n=3 megakaryocyte samples from BLUEPRINT^57^. 52 plasma samples from 28 SARS-CoV-2 patients^42^ were downloaded as FASTQ files were processed as described above. Because of the low coverage (1-2X) of these samples, we extended the marker set from top 25 to top 250 markers per cell type. Roadmap Epigenomics samples^5^ were processed as described above and analyzed using the UXM algorithm. Predicted cell type contributions were averaged across replicates.

## Data availability

The Human DNA Methylation atlas data are deposited in the Gene Expression Omnibus database, accession GSE186458 with access code cjujgaisfjkxnmz. Code is available at https://github.com/nloyfer/wgbs_tools

## Supplemental Data

**Extended Table S1**. List of 207 samples, including tissue of origin, cell type, cell type group, germinal layer of cell type, source of tissue sample, clinical details of donor, sorting details, and sequencing depth. Abbreviations: Had-HU = Hadassah Hospital/Hebrew University; KI = Karolinska Institute; IIDP = Integrated Islet Distribution Program; OHSU = Oregon Health & Science University; UA = University of Alberta

**Extended Table S2**. List of methylation blocks with ≥4 CpGs that show the highest variability (top 1%) across samples.

**Extended Table S3**. List of 39 cell types (top) and 12 super groups (bottom).

**Extended Table S4**. List of 953 cell type-specific unmethylated markers. Also included are additional 279 markers, uniquely unmethylated in combinations of a few related cell types (e.g. pancreatic alpha, beta, and delta cells, gastrointestinal cell types, etc.).

**Extended Table S5**. Cell type-specific markers are enriched for functional terms. Shown are some of the top 10 terms for each marker group, with a significance threshold of FDR≤1E-10.

**Extended Figure S2.**
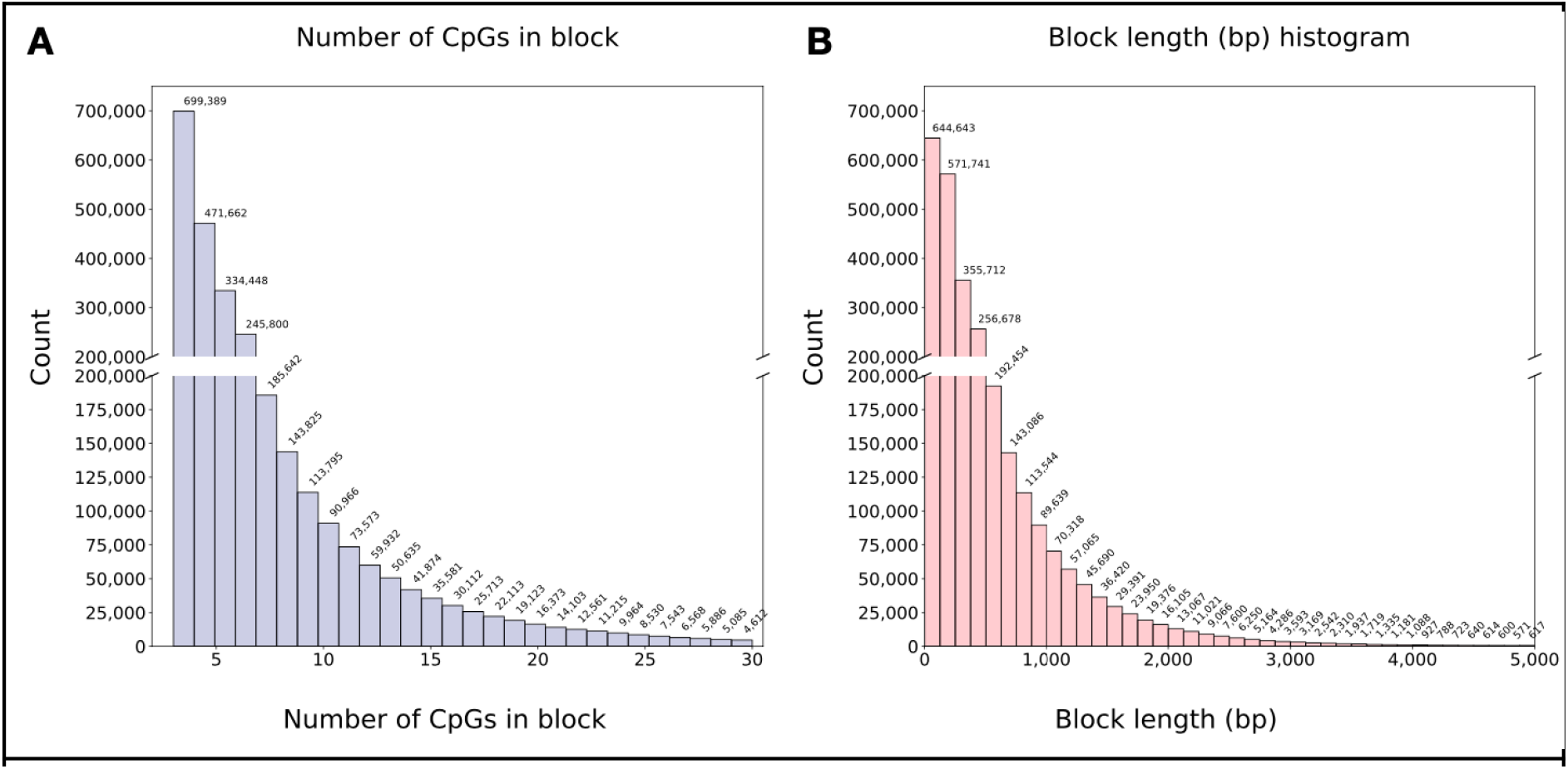
Segmentation of the human genome into 7,264,350 continuous homogeneous blocks. The histograms show the number of segmented blocks as a function of the number of CpGs they contain (left), or their length in bases (right), or as a function. In addition to the 2,746,623 blocks of length 3-30 CpGs (plotted above), there were additional 3,271,607 blocks of one CpG, and 1,185,719 blocks of two CpGs, as well 60,401 of >30 CpGs.

**Extended Figure S3.**
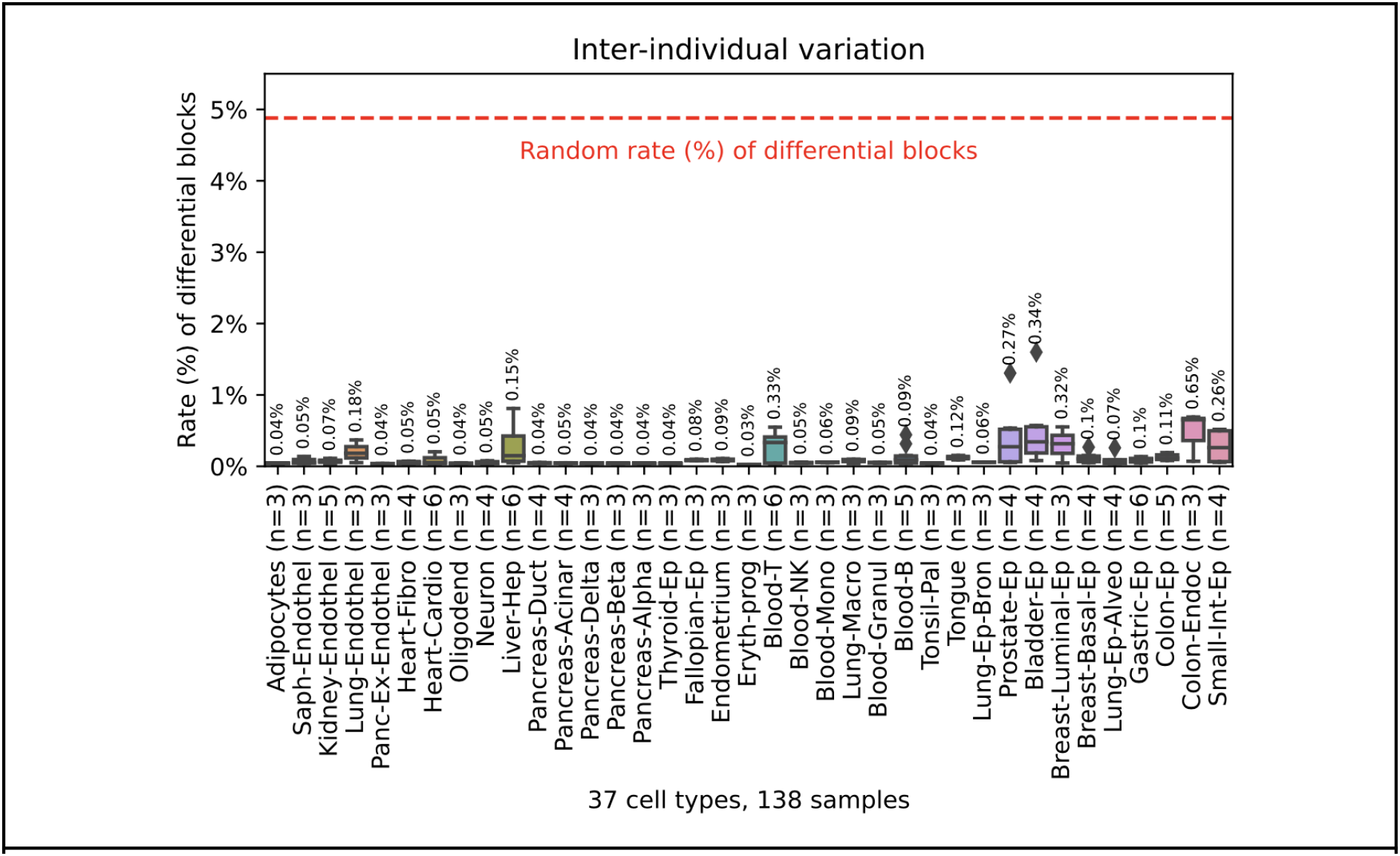
Biological replicates of the same cell type, from different individuals show a surprisingly low rate of differentially methylated blocks. We focused on 37 cellular subtypes with n≥3 replicates (e.g. endothelial cells from a specific tissue) and measured the average percent of methylation blocks (≥3 CpGs) that differ in their methylation by 50% (absolute delta beta), across replicates (shown as Y-axis). Nearly all cellular subtypes (36/37) differ by ≤0.5% of blocks suggesting a very high degree of conservation among replicates. Dotted red line marks the average number of differential blocks between two random samples of different cell types (4.9%).

**Extended Figure S4:**
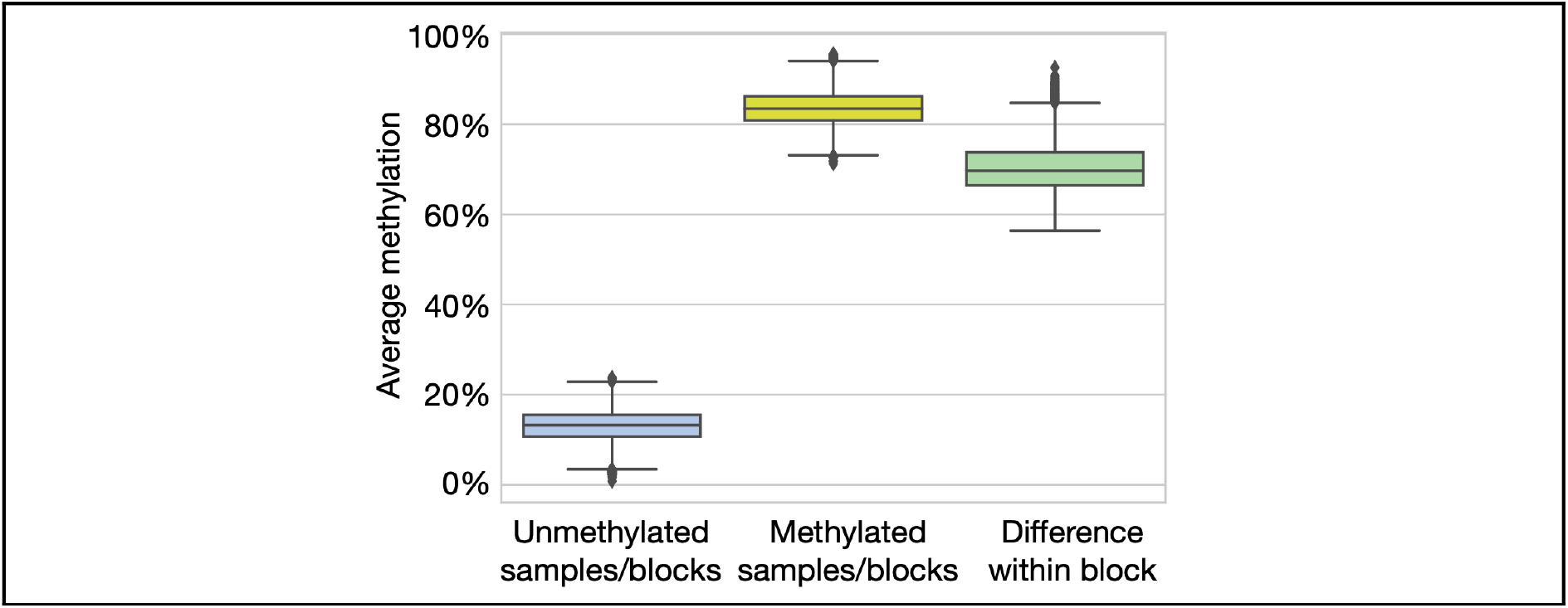
Average methylation in top differentially methylated blocks. Shown are the average methylation values at the 1% most variable blocks of 4 CpGs or more (21,077 blocks). For each block, we computed the average methylation in each sample, and classified them as unmethylated (<50%) or methylated (>50%). Boxplots show the 25^th^ through 75^th^ percentiles among the average methylation levels in unmethylated blocks/samples (blue), methylated ones (yellow) or the difference between methylated and unmethylated samples in the same block (green).

**Extended Figure S5.**
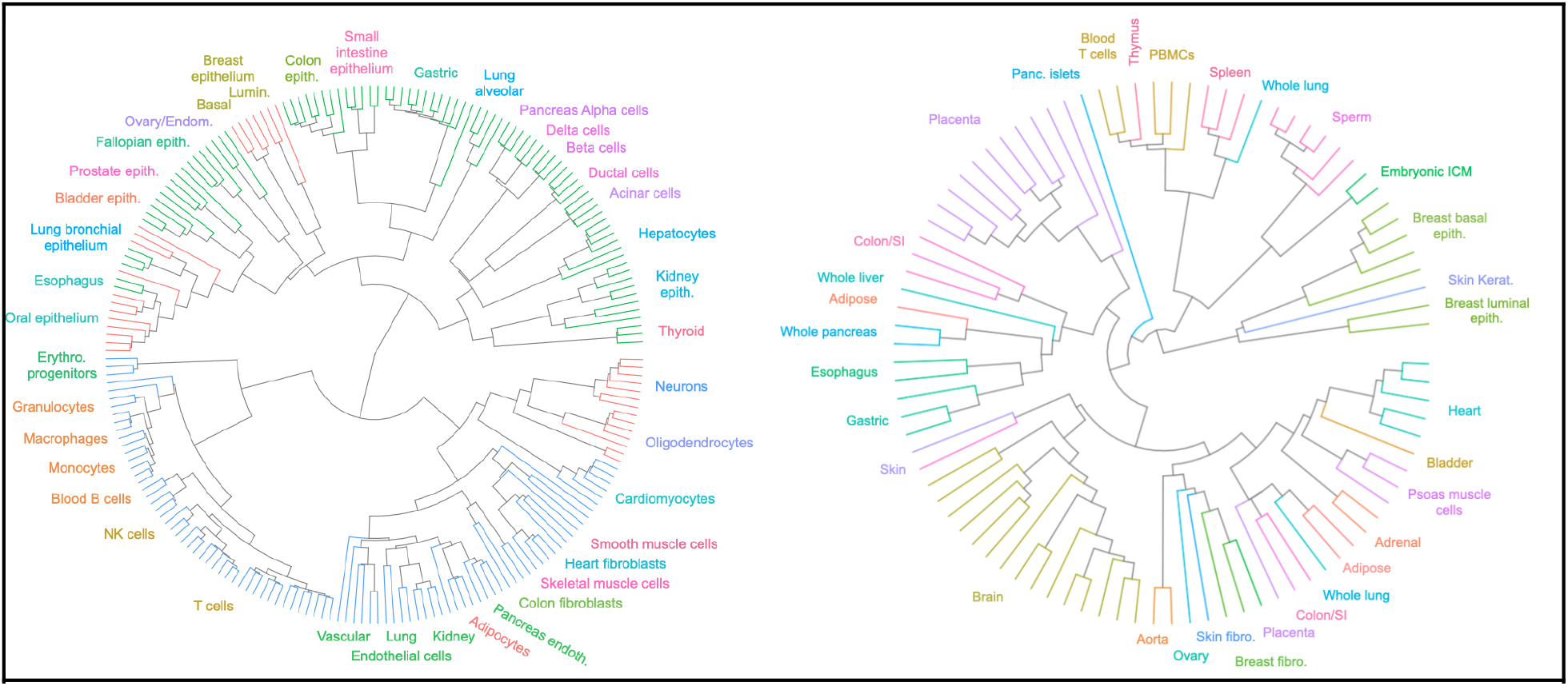
(A) Same as Figure 2, colored by developmental lineage from germ layers, including endoderm (green), mesoderm (blue), and ectoderm (red). (B) same as Figure 2, for Roadmap Epigenomics DNA methylation atlas.

**Extended Figure S6.**
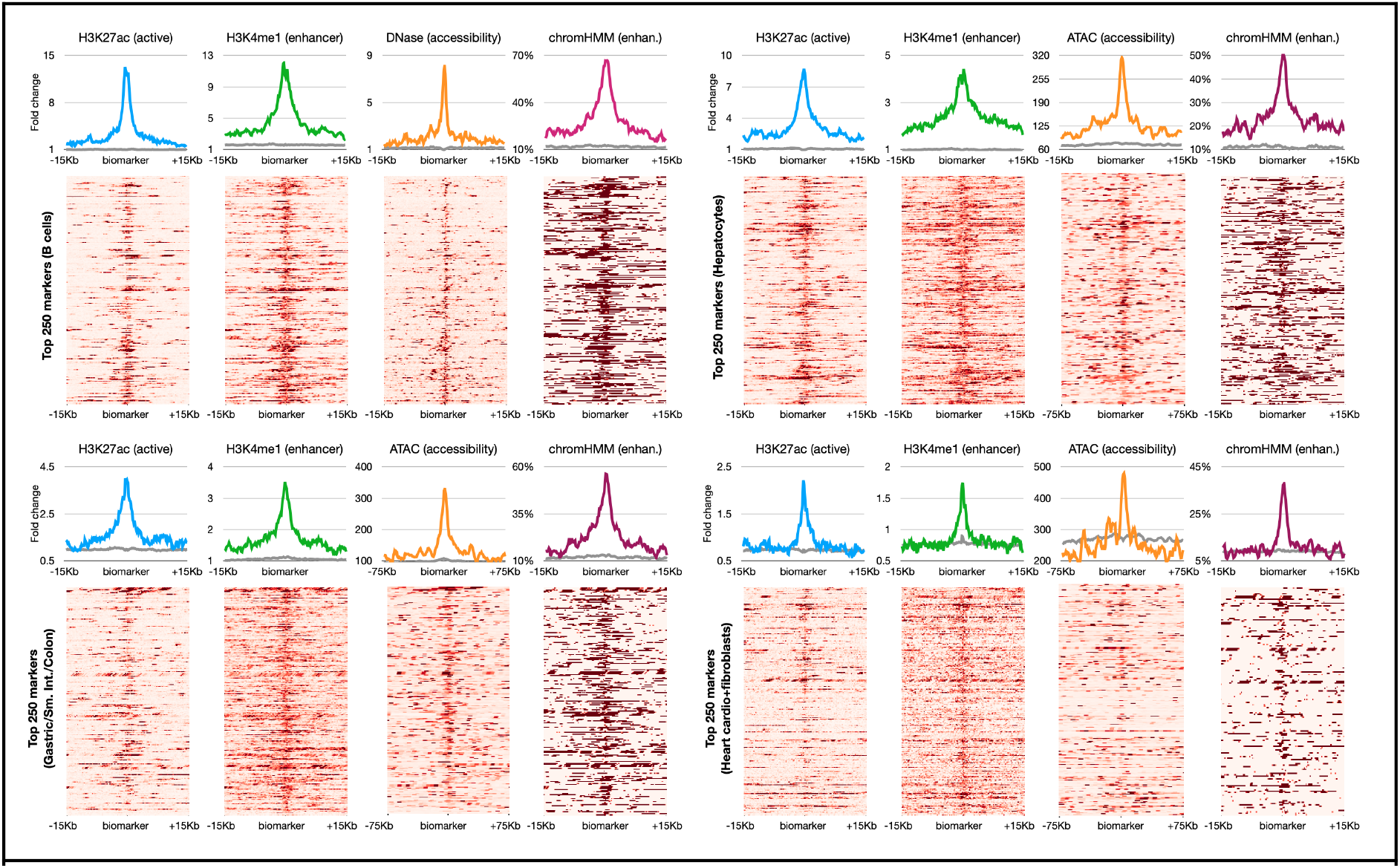
Markers of putative enhancers in other atlas cell types, including top 250 unmethylated markers for B cells (top left), Hepatocytes (top right), Gastric/Small Intestine/Colon epithelium (bottom left), and cardiomyocytes/heart fibroblasts (bottom right). Gray lines mark the same ChIP-seq/ATAC/DNase/chromHMM signal, averaged across all 11,371 unmethylated markers (top 250 per cell type).

**Extended Table S6. Transcription factor binding site enrichment among top 1000 uniquely unmethylated regions for each cell type.**

**Extended Table S7.**
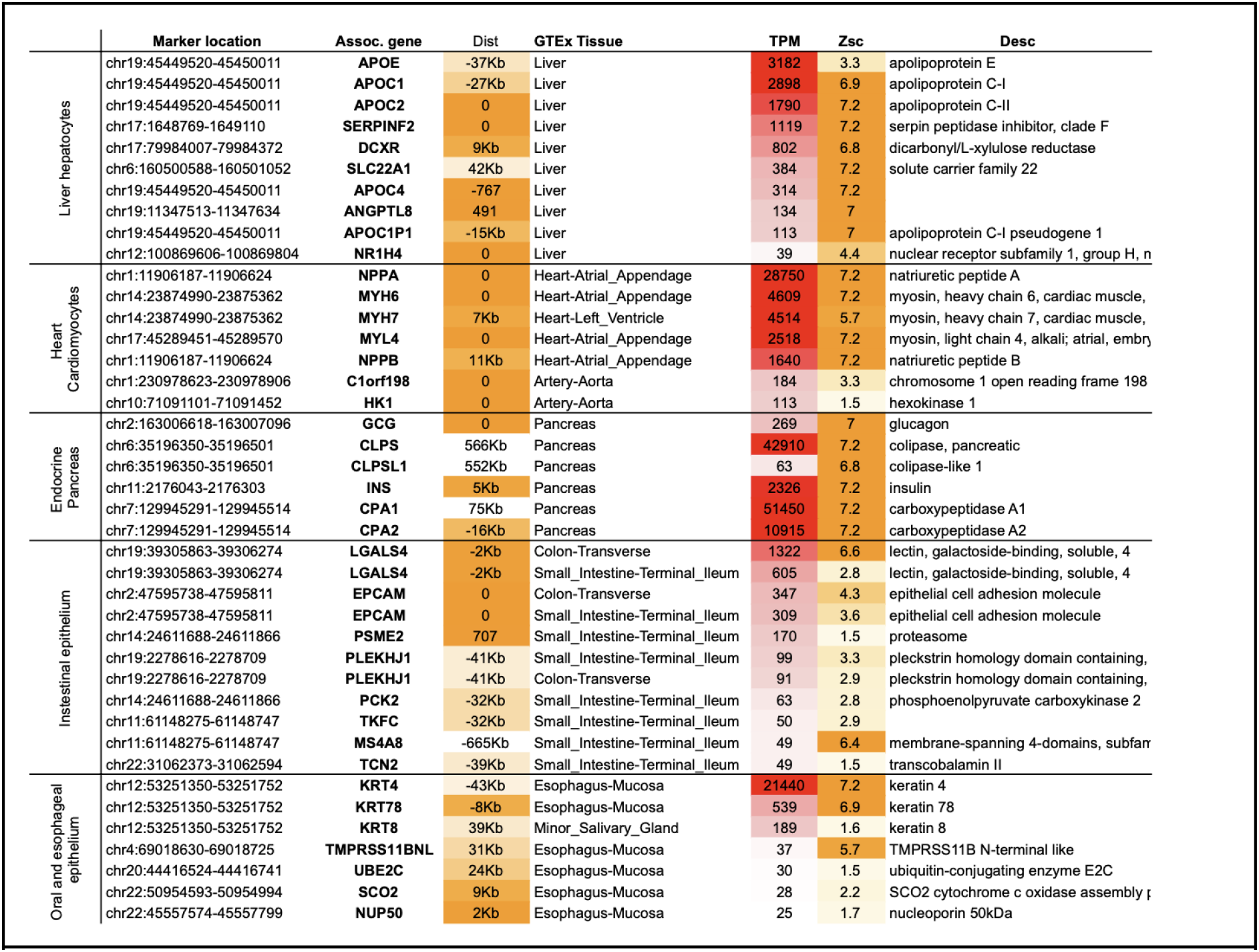
Methylation marker-gene associations. Each cell type-specific unmethylated region was associated with over-expressed genes (up to 750Kb) in related tissues.

**Extended Table S8. Fragment-level deconvolution of 23 blood and 23 plasma samples.**

**Extended Table S9. Fragment-level deconvolution of 104 WGBS samples from Roadmap and ENCODE.**

## Notes

### Competing Interest Statement

This work was supported by GRAIL, Inc. (Menlo Park, CA). GC, JB, HA, PM, PN, AA, OV, and AJ are employees, shareholders and/or founders at GRAIL, Inc. The remaining authors have declared no conflict of interest.

